# Evaluation of Bayesian Linear Regression Models for Gene Set Prioritization in Complex Diseases

**DOI:** 10.1101/2024.02.23.581718

**Authors:** Tahereh Gholipourshahraki, Zhonghao Bai, Merina Shrestha, Astrid Hjelholt, Mads Kjølby, Palle Duun Rohde, Peter Sørensen

## Abstract

Genome-wide association studies (GWAS) provide valuable insights into the genetic architecture of complex traits, yet interpreting their results remains challenging due to the polygenic nature of most traits. Gene set analysis offers a solution by aggregating genetic variants into biologically relevant pathways, enhancing the detection of coordinated effects across multiple genes. In this study, we present and evaluate a gene set prioritization approach utilizing Bayesian Linear Regression (BLR) models to uncover shared genetic components among different phenotypes and facilitate biological interpretation. Through extensive simulations and analyses of real traits, we demonstrate the efficacy of the BLR model in prioritizing pathways for complex traits. Simulation studies reveal insights into the model’s performance under various scenarios, highlighting the impact of factors such as the number of causal genes, proportions of causal variants, heritability, and disease prevalence. Application of both single-trait and multi-trait BLR models to real data, specifically GWAS summary data for type 2 diabetes (T2D) and related phenotypes, identifies significant associations with T2D-related pathways. Furthermore, comparison between single- and multi-trait BLR analyses highlights the superior performance of the multi-trait approach in identifying associated pathways, showcasing increased statistical power when analyzing multiple traits jointly. Additionally, enrichment analysis with integrated data from various public resources supports our results, confirming significant enrichment of diabetes-related genes within the top T2D pathways resulting from the multi-trait analysis. The BLR model’s ability to handle diverse genomic features, perform regularization, conduct variable selection, and integrate information from multiple traits, genders, and ancestries demonstrates its utility in understanding the genetic architecture of complex traits. Our study provides insights into the potential of the BLR model to prioritize gene sets, offering a flexible framework applicable to various datasets. This model presents opportunities for advancing personalized medicine by exploring the genetic underpinnings of multifactorial traits, potentially leading to tailored therapeutic interventions.

## INTRODUCTION

Complex diseases, such as Type 2 Diabetes (T2D), are under the influence of both genetic and environmental (such as socioeconomic and lifestyle) factors (1, 2). Understanding the complex relationship between genetic variation and disease susceptibility is a crucial area of research in genomics. Identification of single genetic variants (commonly known as single nucleotide polymorphisms [SNPs]) associated with phenotypic variation is obtained through genome-wide association studies (GWAS) (3). While GWASs have played a significant role in identifying individual genetic loci associated with disease, they may not fully capture the collective influence of functionally related genes within biological pathways. To address this limitation, gene set analysis has emerged as a valuable analytical tool that focuses on the coordinated action of genes within predefined gene sets (4). The basic idea is to assess whether sets of genes that share common biological functions, such as molecular pathways, display statistical association with the trait or disease.

Biological pathways are complex, interconnected series of molecular actions, genetically encoded within the genome, that regulate various cellular physiological and biochemical processes. Genetic variants associated with complex diseases, such as cancer, metabolic, neurological, and immune-related diseases, tend to be enriched in biological pathways (5, 6). Genetic analyses of biological pathways play a central role in understanding the etiology of complex diseases and hold great potential to identify novel drug targets through elucidating unknown disease mechanisms (6–13).

During the last decade, many different gene set analysis approaches have been proposed (4), including MAGMA (Multi-marker Analysis of GenoMic Annotation) (14), which has become one of the standard tools. MAGMA employs a linear regression model to determine the collective association of gene sets with a disease. Initially, SNP-level statistics (GWAS summary data) within each gene are aggregated while considering the number of SNPs and the degree of linkage disequilibrium (LD) to derive gene-level statistics. In the linear model, the gene-level statistics serve as the response variable, while the gene sets (represented in a binary matrix indicating gene membership) are the predictors. The estimated regression coefficients for each gene set indicate the strength of association with the traits. The significance of these coefficients is assessed against a null distribution, typically generated through permutations or a model-based approach, indicating to which extent each gene set is associated with the trait of interest.

Handling many gene sets (such as biological pathways) can introduce several challenges. Firstly, overfitting becomes a concern because the gene set model may fit noise instead of underlying biological signals. Secondly, many gene sets are correlated due to biological interconnectedness, and because all gene sets are fitted jointly in the MAGMA model, multicollinearity becomes an issue. Thirdly, the risk of false positives also escalates with more predictors, necessitating stringent multiple-testing corrections. Lastly, the abundance of gene sets complicates the interpretation of results, making it challenging to discern the individual contributions of each set to the phenotype. Here, we propose a strategy to address these issues by implementing variable selection and regularization within the MAGMA framework to enhance model robustness and interpretability.

The Bayesian Linear Regression (BLR) model is one of the procedures that can overcome the limitations of the standard linear model used in the MAGMA procedure. The Bayesian framework effectively handles multiple testing issues, reducing the risk of false positives, which is common when testing numerous gene sets. Additionally, it addresses the challenge of gene set overlap and interdependency. The use of spike-and-slab priors aids in variable selection and regularization by better distinguishing between true associated gene sets from those that are significant because of partially shared genes. Because the BLR framework is flexible, it allows multiple traits to be analyzed jointly. Thus, incorporating correlated trait information in gene set analysis provides deeper insights by identifying shared genetic factors, further enhancing our understanding of complex biological processes (4, 14–16).

The aim of this study was to present and evaluate a gene set prioritization approach utilizing BLR models within the MAGMA gene-set analysis procedure. To investigate how different characteristics of gene sets and different trait genetic architectures influenced the detection power, we conducted a comprehensive simulation study to assess the model’s statistical performance utilizing genetic data from the UK Biobank (17). We subsequently applied our BLR prioritization methodology to publicly available GWAS summary data for nine distinct complex traits. To uncover the shared genetic architecture among these traits, we advanced our analysis by developing a multi-trait BLR model. This enhancement allowed for the simultaneous integration of GWAS information across all nine traits, facilitating a more comprehensive analysis.

## MATERIAL AND METHOD

Figure 1 presents a schematic overview of the workflow. In the initial step, GWAS summary data for the traits of interest are utilized to compute gene-level Z-scores using the VEGAS (Versatile Gene-Based Association Study) approach (18). We constructed a design matrix linking genes to gene sets to integrate curated gene sets. The BLR model was then fitted using this design matrix of all gene sets as input features (predictors) and the Z-scores as the response variable. This results in a posterior inclusion probability (PIP) for each gene set, which represents the probability that the gene set is included in the model. Gene sets with higher PIPs are given higher priority scores, facilitating the identification of potential biological mechanisms underlying the observed genetic associations. Notably, our methodology extends to a multiple-trait analysis, enabling a comprehensive exploration of gene sets across diverse traits. Details on the statistical model and analyses, VEGAS approach, and used data are provided in the subsequent sections.

**Fig. 1.**
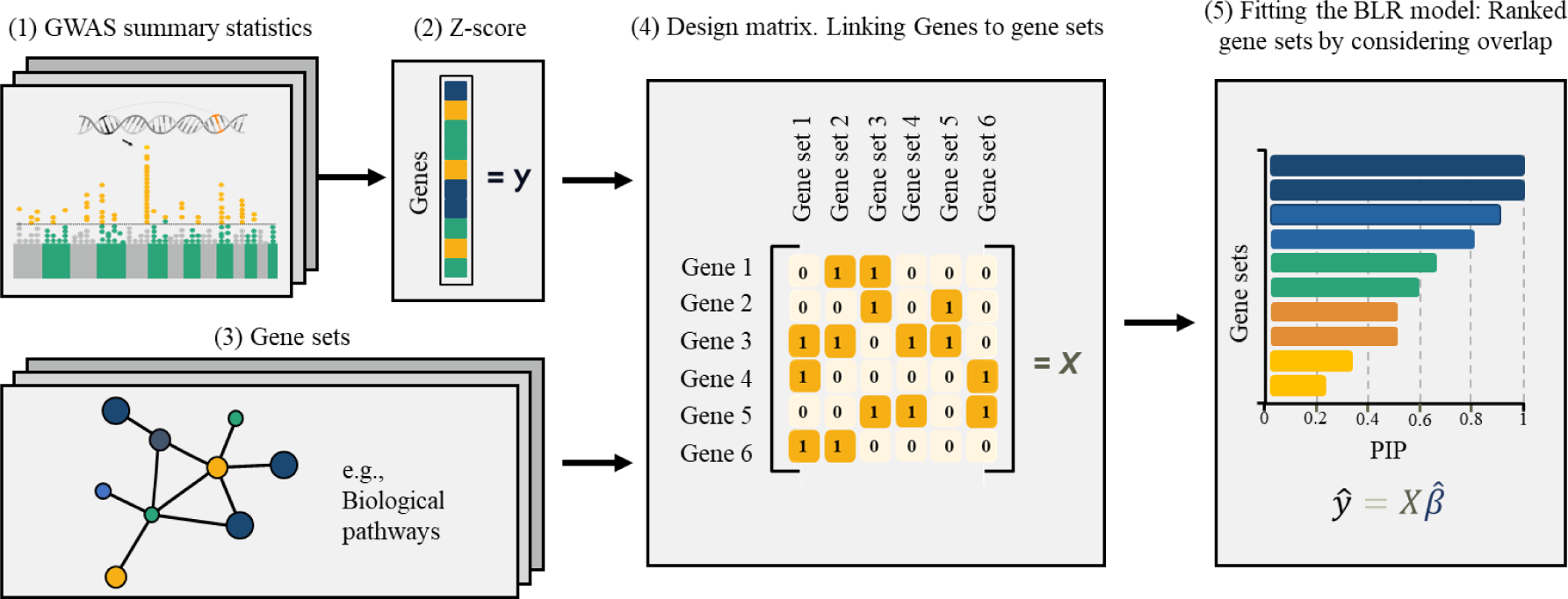
Overview of gene set prioritization method using BLR model. GWAS: Genome-Wide Association Study. PIP: Posterior Inclusion Probability.

### Statistical models and analyses

#### Linear model for gene set analysis

The foundation of our approach rests upon a linear model that can be expressed in matrix notation as follows:

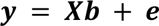

where ***y*** represents the per-gene statistic, such as the gene-level Z-score (see section 2.1.5), indicating the strength of association between individual genes and the trait phenotype, ***X*** is a design matrix linking genes to gene sets, as well as the corresponding per-gene statistic, and ***e*** denotes the residuals, which are assumed to follow an independent and identically distributed normal distribution with a mean of 0 and variance σ^2^. The dimensions of ***y***, ***X***, ***X*** and ***e*** depend on the number of traits (*k*), the number of gene sets (*m*), and the number of genes (*n*). The design matrix ***X*** has the dimension *n*-by-*m*, which takes the value one if a gene belongs to a gene set; otherwise, the elements are zero. The vector ***X*** represents the regression coefficient for each gene set.

#### Single trait BLR model

We used a BLR model using a BayesC (19) prior assumptions to model the association between gene sets and traits. BayesC utilizes a spike-and-slap prior distribution:

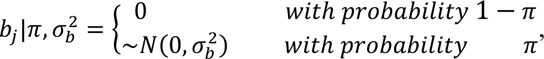

assuming the regression effects (***X***) are drawn from a mixture distribution comprising a point mass at zero and a normal distribution defined by a common variance 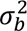 for the regression effects. Each regression effect (*b*_*j*_) is either zero or non-zero, where zero implies insignificance, and non-zero signifies a contribution to the response variable. The prior probability, π = 0.001, determines the proportion of regression effects falling into either class. The prior distribution of the common variance 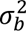 for the regression effects follows an inverse Chi-square distribution, χ^−1^(*S*_*b*_, *v*_*b*_), where *S*_*b*_ represents the scale parameter of an inverse Chi-square distribution and *v*_*b*_ represents the degrees of freedom parameter.

The mixture proportions are determined using a Dirichlet distribution (*C*, *c* + α), where *C* represents the number of mixture components in the distribution of regression effects, *c* represents the vector of counts of regression variables within each component, and α = (1,1). To manage these complex distributions and to facilitate the analysis, a variable called *d* = (*d*_1_, *d*_2_ …, *d*_*m*−1_, *d*_*m*_) is added using the idea of data augmentation, and it shows whether the *j*^*th*^ regression effect is zero or nonzero.

#### Multiple-trait BLR models

BLR models can be extended to encompass multiple traits, which is useful for identifying common biological functions or gene sets shared across traits or diseases. We implemented a general multiple-trait Bayesian linear regression model based on the BayesC prior (19). This model enables a gene set to influence any combination of traits, offering insights into whether gene sets affect all, some, or none of the traits. The multiple-trait BLR model is subject to regularization, similar to the single-trait model, while leveraging information from correlated traits. The core equation governing the multiple-trait BLR model is represented as:

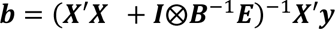

In this model, the key parameters include the covariance matrix for the regression effects, denoted as ***B***, and the residual covariance matrix, denoted as ***E***. These matrices capture the shared relationships between regression effects across traits.

#### Implementation of BLR model analysis

The BLR model parameter estimates 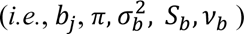 were obtained using Markov Chain Monte Carlo (MCMC) sampling procedures as implemented in the blr function in the qgg package. Further details on these procedures are provided in the Supplementary Note and by Rohde et al. (16). For analysis involving both single-trait and multiple-trait scenarios, a total of 3000 iterations were employed, with the initial 500 iterations designated as burn-in to ensure adequate model convergence. Multiple runs were conducted to confirm convergence.

#### Gene-level statistics

Gene-level association statistics (***Z***) were computed using the VEGAS (versatile gene-based association study) approach(18), as 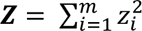 where *z* is the standardized single marker GWAS regression coefficients 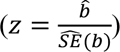, while accounting for the LD between markers in the genes. Gene-based *P*-values was computed based on the distribution of quadratic forms in normal variables using saddle point approximations (20, 21) as implemented in the vegas function in the qgg package. To perform the gene-set analysis, for each gene g the gene *P*-value *p*_*g*_ computed with the gene analysis is converted to a Z-value *z*_*g*_ = Φ^−1^(1 − *p*_*g*_), where Φ^−1^ is the probit function. This yields a roughly normally distributed variable Z with elements *p*_*g*_that reflects the strength of the association each gene has with the phenotype, with higher values corresponding to stronger associations. Ancestry matched LD (i.e., European) information for each gene region was obtained from the 1000 Genomes Project reference panel (22),

### Simulation study

#### Simulation of phenotypes

The primary aim of the study was to evaluate the BLR gene-set prioritization approach, which we assessed using comprehensive simulations. Genetic variants originating from UKB chip genotypes were used to simulate quantitative and binary traits restricting to unrelated individuals of White British origin (*n*=335,532). Initial quality controls of genetic variants were performed such that SNPs with a minor allele frequency below 0.01, a genotype call rate lower than 0.95, and those not conforming to Hardy-Weinberg equilibrium (with a *P*-value of 1 × 10^−12^) were excluded. Additionally, genetic variants within the major histocompatibility complex, exhibiting ambiguous alleles (such as GC or AT), having multiple alleles, or representing indels, were removed (23), yielding a final set of 533,679 SNPs.

Various simulation scenarios were explored, considering factors such as trait heritability (ℎ^2^ = 0.1 *pp* 0.3), the proportion of causal genetic variants (π = 0.01 *pp* 0.001), and disease prevalence of binary traits (*p* = 0.05 *pp* 0.15). Furthermore, we also considered two different genetic architecture scenarios: GA1 represents a simplified genetic architecture characterized by a mixture of point mass at zero and a single normal distribution of genetic effects. GA2 represents a more complex genetic architecture involving a mixture of multiple normal distributions for genetic effects. In total, this resulted in eight different simulation scenarios for quantitative traits and 16 scenarios for binary phenotypes, with ten replicates for each scenario. Detailed information on the quantitative and binary phenotype simulations is described in the Supplementary Note.

#### Simulation of gene sets

To assess the accuracy of the BLR gene-set prioritization approach, synthetic genes and gene sets were derived from the simulated phenotype data described above based on the predefined sets of causal SNPs. Initially, genes were divided into two groups: causal genes (*i.e.*, genes containing causal SNPs), and non-causal genes (genes lacking causal SNPs). Two key parameters were used to control the size and enrichment of causal genes within gene sets: the total number of genes in each gene set (referred to as the gene set size), ranging from 10 to 200 genes, and the number of causal genes selected from causal genes. Various values for number of causal genes were explored, including 0, 5, 10, 25, 50, 100, and 200. Ten replicates were performed for each gene set configuration to address sampling bias. In total, 21 distinct gene set configurations were generated for each simulation scenario. Scenarios with gene sets containing no causal genes were defined as controlled configurations. Detailed scenarios for quantitative and binary phenotypes are outlined in Table 1.

**Table 1.** Simulated phenotype scenarios (Binary and quantitative traits)

| $h^2$ | $\pi$ | GA | Quantitative phenotype scenarios | P(binary traits) | Binary phenotype scenarios |
| --- | --- | --- | --- | --- | --- |
| 0,3 | 0,001 | GA1 | Sim1 | 0,05 | sim1 |
|  |  |  |  | 0,15 | sim2 |
| 0,3 | 0,001 | GA2 | Sim2 | 0,05 | sim3 |
|  |  |  |  | 0,15 | sim4 |
| 0,3 | 0,01 | GA1 | Sim3 | 0,05 | sim5 |
|  |  |  |  | 0,15 | sim6 |
| 0,3 | 0,01 | GA2 | Sim4 | 0,05 | sim7 |
|  |  |  |  | 0,15 | sim8 |
| 0,1 | 0,001 | GA1 | Sim5 | 0,05 | sim9 |
|  |  |  |  | 0,15 | sim10 |
| 0,1 | 0,001 | GA2 | Sim6 | 0,05 | sim11 |
|  |  |  |  | 0,15 | sim12 |
| 0,1 | 0,01 | GA1 | Sim7 | 0,05 | sim13 |
|  |  |  |  | 0,15 | sim14 |
| 0,1 | 0,01 | GA2 | Sim8 | 0,05 | sim15 |
|  |  |  |  | **0,15** | **sim16** |

$\pi$ : proportion of causal markers, $h^2$ : heritability, GA: genetic architecture, P: prevalence

#### Single marker regression analysis of simulated data

Standard GWASs were conducted for each simulated phenotype, splitting the data into five cross validation replicates, each comprising training (80%) and validation (20%) subsets. The GWAS procedure was separately performed in the training populations for each of the five replicates. For quantitative phenotypes, we utilized single-marker linear regression models with the R package qgg (16, 24), and for binary phenotypes, single-marker logistic regressions were conducted using PLINK 1.9 (25).

#### Evaluation metrics of simulation study

To assess the accuracy of the BLR model in gene set prioritization for the simulated data, we utilized the F1 classification score as a key performance metric. The F_1_ score ranges from 0 to 1 and combines precision (*p*) and recall (*p*) to provide a balanced assessment of the performance of the model. Values close to 1 refer to the capability of the BLR model to identify true associated gene sets better and reduce false positives. It is expressed as:

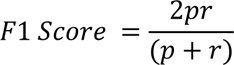

Precision (*p*) measures the accuracy of identifying relevant gene sets among those predicted as significant, computed as *p* = *p = TP + FP*), where *TP* is true positives (correctly identified relevant gene sets) and *FP* is false positives (incorrectly identified gene sets). Recall (*p*) evaluates the model’s ability to correctly identify truly relevant gene sets and is calculated as *p* = *TP*⁄(*TP* + *FN*), with *FN* representing false negatives (relevant gene sets missed by the model) (26).

### Data processing and integration

Data processing and integration were facilitated by using the R **gact** package, which is designed for establishing and populating a comprehensive database focused on genomic associations with complex traits. The package has two primary functions: infrastructure creation and data acquisition. It facilitates the assembly of a structured repository that includes single marker associations, all rigorously curated to ensure high-quality data. Beyond individual genetic markers, the package integrates a broad spectrum of genomic entities, encompassing genes, proteins, and a variety of biological complexes (chemical and protein), as well as various biological gene sets. Details of this package, including examples of analysis scripts used for analysing real traits in this study, can be found can be found in the package documentation (27).

#### GWAS summary data

We applied the BLR models to nine distinct traits with publicly available GWAS summary data. These include Type 2 Diabetes (T2D) (28), Coronary Artery Disease (CAD) (29), Chronic Kidney Disease (CKD) (30), Hypertension (HTN) (31), Body Mass Index (BMI) and Waist-Hip Ratio (WHR) (32), Glycated Hemoglobin (Hb1Ac) (30), Height (33), Systolic Blood Pressure (SBP) (34), and Triglycerides (TG) (35). Detailed study information can be found in Supplementary Table S1.

#### Gene annotation and linkage disequilibrium reference data

For the gene-level association statistics using the VEGAS approach, reference data from the 1000 Genomes Project were utilized. The datasets encompass genetic variation across three major populations: European (EUR), East Asian (EAS), and South Asian (SAS).. Initial quality control of genetic variants was performed such that genetic variants with a minor allele frequency below 0.01, a call rate lower than 0.95, and those not conforming to Hardy-Weinberg equilibrium (with a *P*-value of 1 × 10^−12^) were excluded. Additionally, genetic variants situated within the major histocompatibility complex, exhibiting ambiguous alleles (such as GC or AT), having multiple alleles, or representing indels, were removed (23).

Genetic markers located with 35kb upstream and 10kb downstream of the open reading frame were used as the marker set for the gene.

#### Gene sets

Gene sets were derived from a number of different annotation sources. Biological pathways utilized in our study were curated from the Kyoto Encyclopedia of Genes and Genomes (KEGG) (36), a well-established and comprehensive resource for understanding cellular functions and biological processes. KEGG pathways were obtained using the msigdb R package (37). Gene-disease association data were used to enhance our analysis, focusing on comprehensive text-mining results, expert-curated knowledge, experimental evidence, and integrated datasets pertaining to human diseases. The data used included full and filtered datasets from text mining (human_disease_textmining_full.tsv and human_disease_textmining_filtered.tsv), curated knowledge datasets (human_disease_knowledge_full.tsv and human_disease_knowledge_filtered.tsv), experimental datasets (human_disease_experiments_full.tsv and human_disease_experiments_filtered.tsv), and an integrated dataset combining all sources (human_disease_integrated_full.tsv). All files were retrieved from JensenLab (38).

#### Measuring the degree of enrichment

Gene set prioritization was quantified using the PIP. Gene sets with a PIP ≥0.1 in at least one trait were considered associated. Additionally, we utilized another association metric from the BLR model: the posterior mean of regression effects. Negative regression effect values indicated gene sets enriched for non-associated genes, which were excluded to refine our focus on gene sets enriched for associated genes.

#### Enrichment analysis using hypergeometric test

In order to validate that the top-ranking gene sets identified with our BLR method are supported by external evidence, we performed an enrichment analysis using a hypergeometric test. For every gene set, we tested for enrichment of disease-gene association obtained from the DISEASES database (39, 40), which provides disease–gene association scores derived from curated knowledge databases, experiments primarily GWAS catalog, and automated text mining of the biomedical literature. The enrichment analyses were conducted on integrated and individual channels, including knowledge base, text mining, and experiment.

## RESULTS

### Simulation study

The simulation study aimed to assess the performance of the BLR model to prioritize gene sets for their association with a phenotype. By examining various trait and gene set characteristics, the objective was to understand the model’s behavior and its ability to handle challenges inherent in real data applications.

#### Effect of gene set characteristics

We evaluated the performance of the BLR model by considering various factors, including the size of the gene set (i.e., the number of SNPs), the number of causal genes, and the proportion of causal genes within the gene set. Increasing the number of causal genes in the gene set consistently led to an increase in the F1 score across all gene sets of the same size (Figure 2). However, when gene sets contained the same number of causal genes, increasing the total number of genes tended to decrease the F1 score. Additionally, gene sets containing more genes exhibited a larger F1 score when the proportion of causal genes remained constant.

**Fig. 2.**
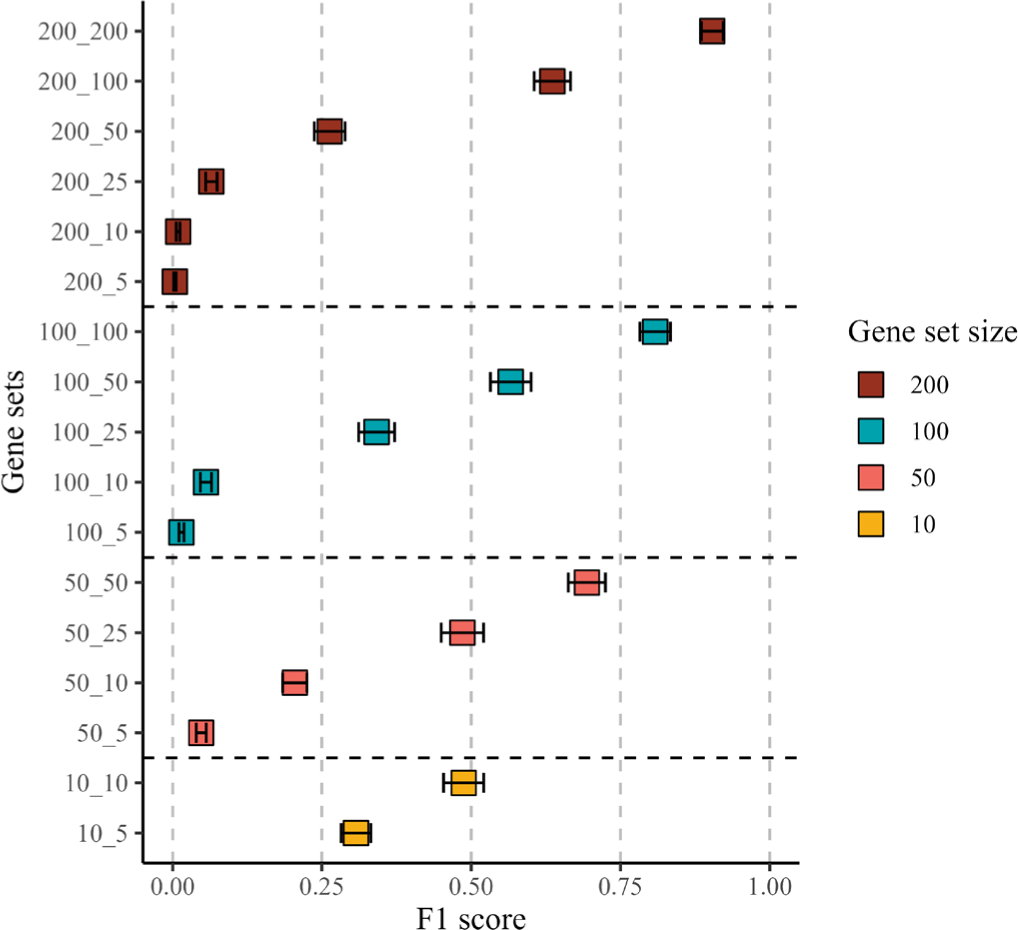
Assessing BLR model performance across gene set configurations in simulated data (binary traits). The y-axis represents pathways, with the first number indicating the size of the pathway and the second number representing the number of causal genes within the pathway. The x-axis displays the mean F1 score across all simulation scenarios. Points represent

#### Effect of trait characteristics

We then investigated how different trait characteristics of binary and quantitative phenotypes affected model performance. Specifically, we investigated the impact of heritability (h²), the proportion of causal markers (π), genetic architecture (GA), and the effect of disease prevalence on the model’s ability to identify gene sets containing causal SNPs. Our findings showed that the scenario with a lower proportion of causal markers (π = 0.001) consistently achieved higher F1 scores across all gene sets (Figure 3A). Similarly, the scenario with higher heritability (h² = 0.3) demonstrated superior F1 scores across most gene sets (Figure 3B). Furthermore, GA1, characterized by a single normal distribution, generally outperformed GA2, which involves a more complex architecture of the regression effects with a mixture of normal distributions (Figure 3C). In addition, the scenario where the disease prevalence was highest (p = 0.15) consistently displayed a superior F1 score compared to the scenario with a lower disease prevalence (p = 0.05, Figure 3D). Similar patterns were observed for the simulated quantitative phenotypes (Supplementary Figure S1). Detailed results across all scenarios can be found in Supplementary Tables S2 and S3.

**Fig. 3.**
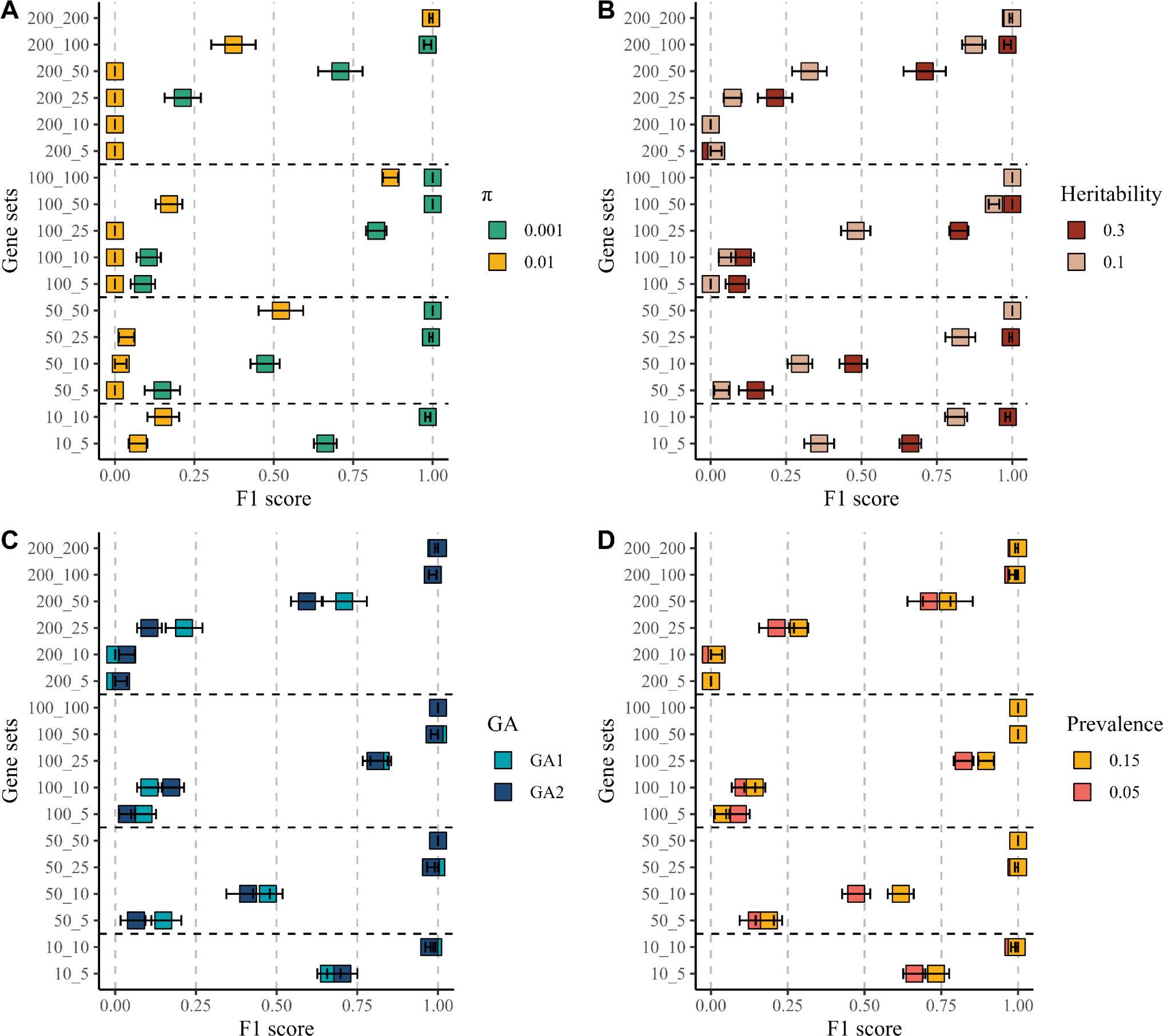
Evaluation of BLR model perfromance in simulation scenarios (binary traits). Scenarios (A-D) were systematically compared by varying a specific property while keeping others constant. **A.** Illustrates the impact of varying the proportion of causal markers (π). **B.** Demonstrates scenarios with varying heritability (h²). **C.** Compares two genetic architecture scenarios, GA1 and GA2. **D.** Highlights the effect of prevalence (p). The y-axis represents pathways, with the first number indicating the size of the pathway and the second number representing the number of causal genes within the pathway. The x-axis displays the F1 score. Points represent mean values across ten replicates, and error bars indicate standard errors.

### Application of BLR model to real data

We employed single and multiple-trait BLR models to investigate gene sets associated with T2D and related phenotypes, utilizing publicly available GWAS summary data. Out of the 186 pathways studied, we identified three KEGG pathways with significant associations with T2D across both models: “*Type II diabetes mellitus*”, “*Type I diabetes mellitus*”, and “*Maturity onset diabetes of the young*” (Figure 4).

**Fig. 4.**
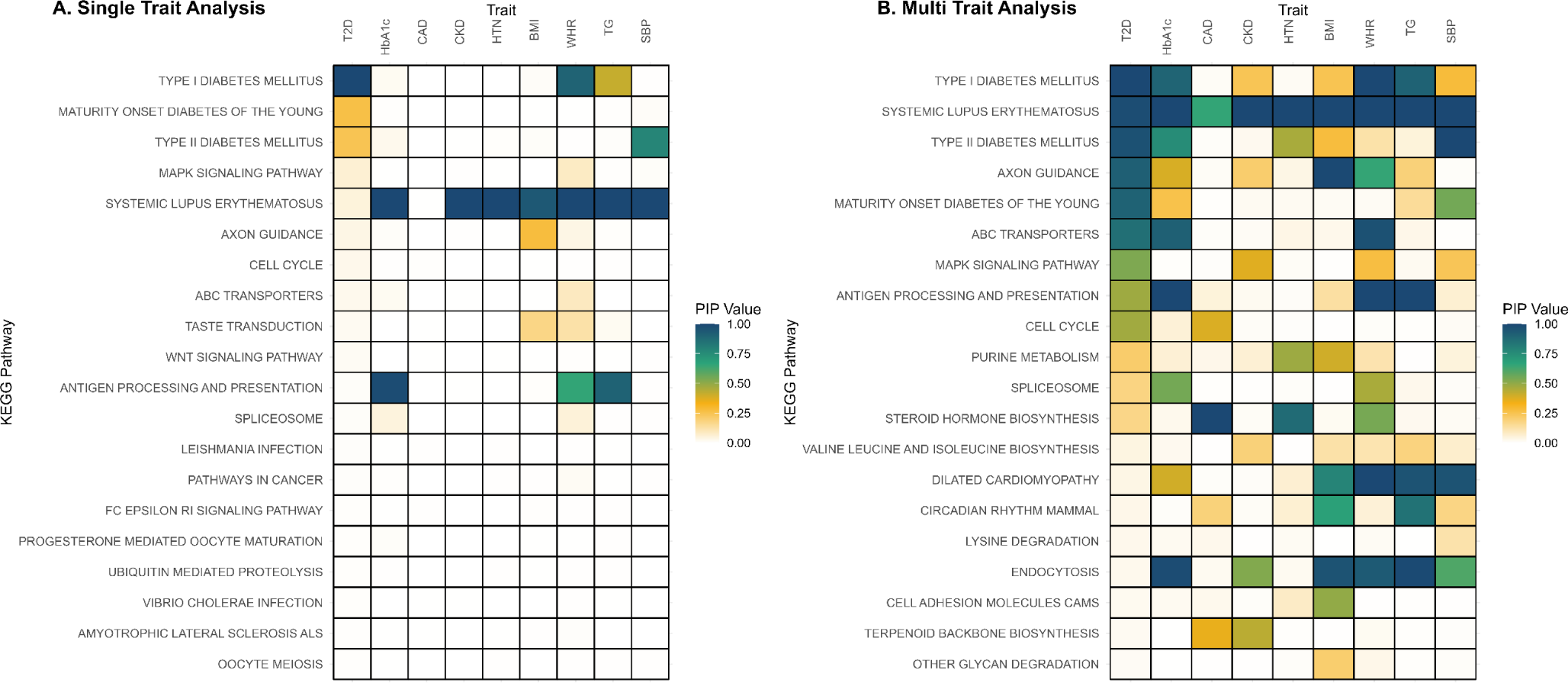
Comparative heatmap analysis of pathway associations with type 2 diabetes and correlated traits using single and multi-trait BLR model. Columns correspond to traits analyzed through GWAS, and rows represent KEGG pathways. Warmer colors indicate stronger associations, as measured by higher Posterior Inclusion Probabilities (PIPs), with a PIP of 1 indicating the highest association level, suggesting a strong likelihood that the pathway is relevant to the trait. Type 2 Diabetes (T2D), Hemoglobin A1c (Hb1Ac), Coronary Artery Disease (CAD), Chronic Kidney Disease (CKD), Hypertension (HTN), Body Mass Index (BMI), Waist-Hip Ratio (WHR), Triglyceride (TG), Systolic Blood Pressure (SBP).

#### Comparison of single-trait and multiple-trait analyses

Utilizing the multiple-trait BLR approach, we found 12 KEGG pathways associated with T2D (Figure 4B), suggesting increased statistical power when jointly analyzing multiple traits. Across all traits, the multi-trait analysis identified more associated pathways and showed higher statistical significance than the single-trait analysis (Figure 4B). Notably, most of the pathways identified as highly associated in the single-trait analysis were also confirmed in the multi-trait analysis (Figure 5). Additional results are available in Supplementary Tables S4 and S5.

**Fig. 5.**
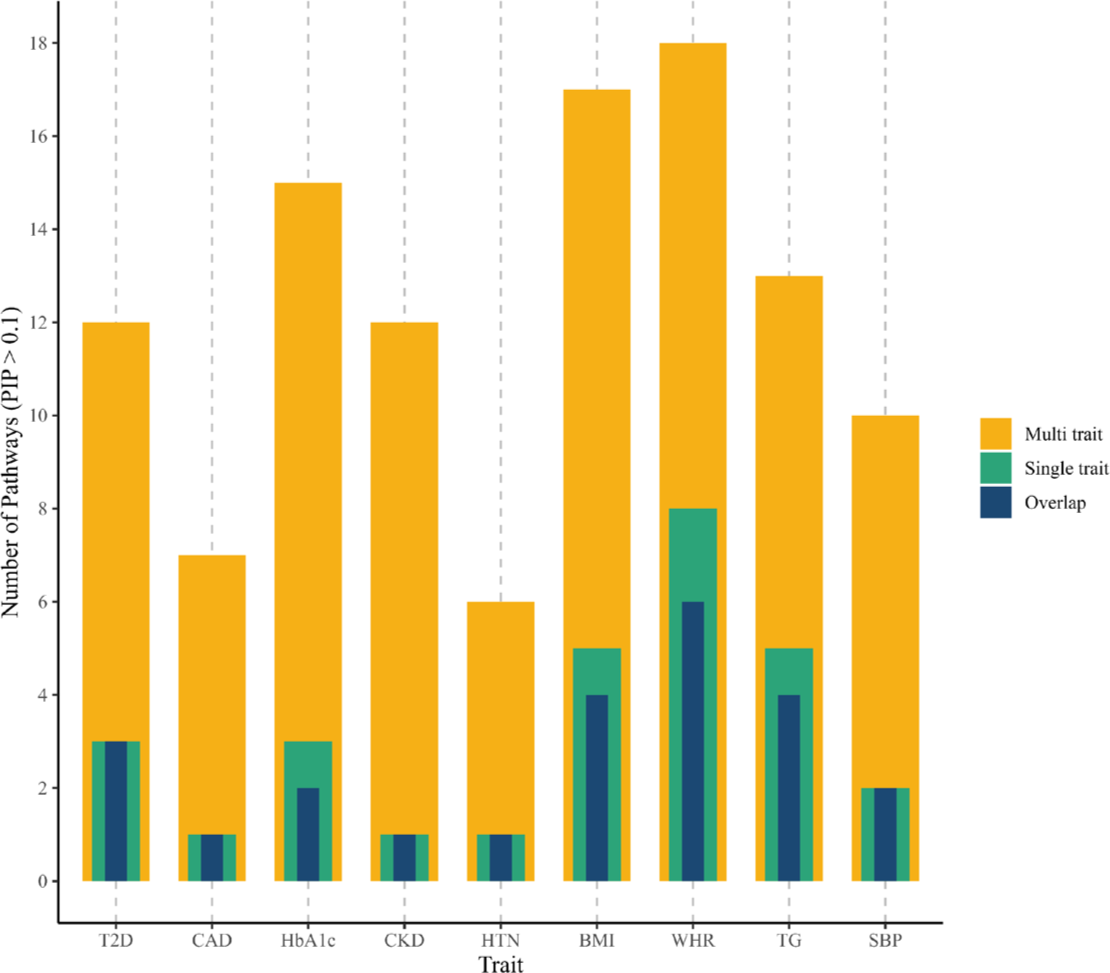
Pathway overlap comparison of single-trait and multi-trait analyses. Numbers represent the total count of pathways with a mean PIP > 0.1.

#### Application of multiple-trait BLR model to different T2D GWAS subgroups

To confirm the robustness and consistency of our findings, we applied the multiple-trait BLR model to distinct T2D GWAS subgroups. Specifically, we investigated gender-based differences by analyzing male and female cohort data. Additionally, we delved into the influence of genetic ancestry by conducting separate analyses for European (EUR), East Asian (EAS), and South Asian (SAS) populations. The highest-ranked pathways within these subgroups exhibited remarkable similarity to the pathways identified in the overall multiple-trait BLR analysis; interestingly, when comparing results between males and females, minimal differences were observed, and the pathway prioritization remained highly consistent across genders (Figure 6A). Similarly, the highest-ranked pathways showed substantial overlap in comparing EUR and EAS ancestries (Figure 6B). However, the results for the SAS subgroup exhibited peculiar patterns. The SAS subgroup analysis may be influenced by a comparatively lower number of individuals in the dataset, potentially contributing to the observed discrepancies.

**Fig. 6.**
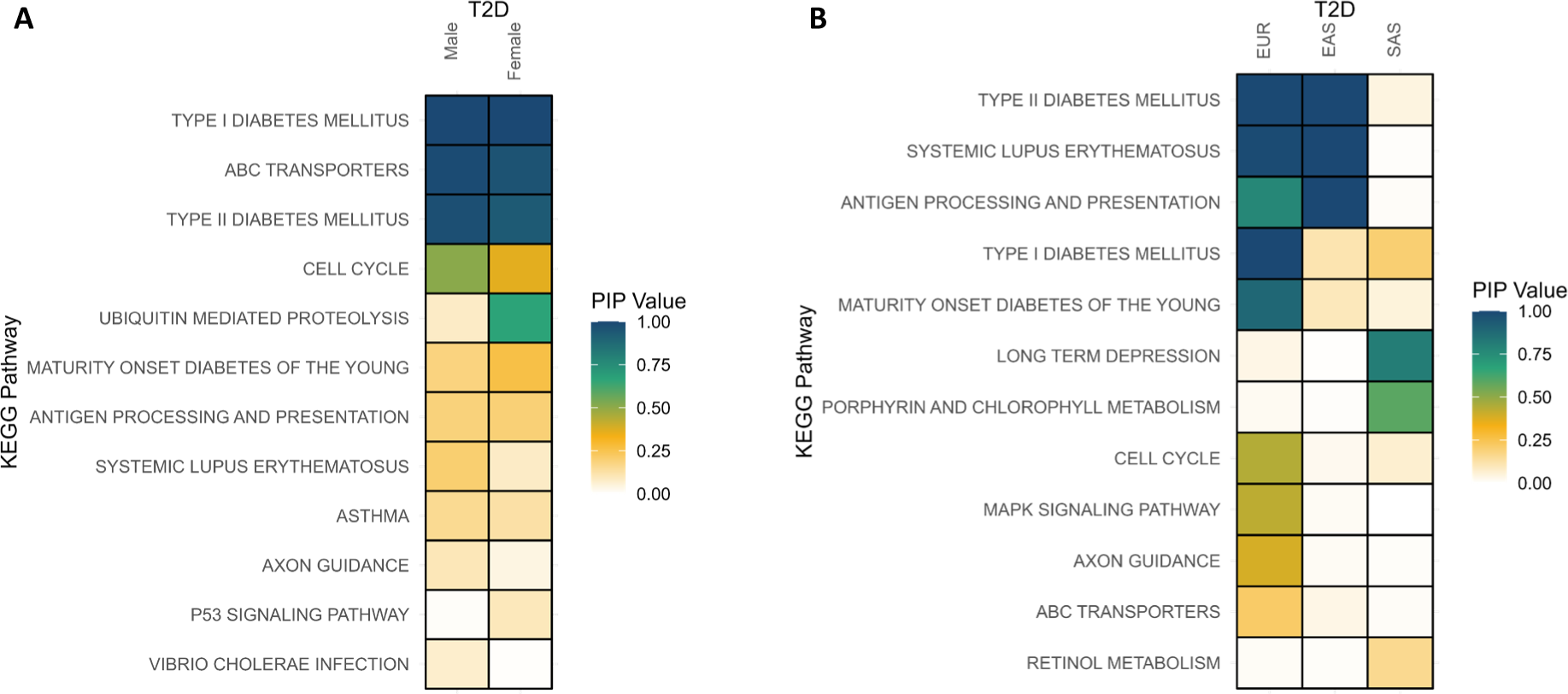
Application of multi-trait BLR model to different T2D GWAS subgroups. Posterior Inclusion Probability (PIP), Type 2 Diabetes (T2D).

#### Analysis of pathway enrichment for T2D

We furthermore conducted a gene set enrichment analysis to explore the relationship between KEGG pathways and diseases, specifically focusing on pathways relevant to diabetes. Utilizing gene-level statistics, we integrated data from various public resources, including text mining (40, 41), experiments (GWAS catalog) (42), and knowledge bases (43) with gene sets representing KEGG pathways. Specifically, we targeted the disease term “diabetes,” excluding other known types such as type 1, maturity-onset, neonatal, and gestational diabetes. Employing a hypergeometric gene set testing approach, we found a significant enrichment of diabetes-related genes within the top T2D pathways resulting from the multiple-trait analysis. We observed that the majority of highly associated pathways exhibited remarkably similar significant *P*-values (Table 2, *P*-value < 0.05). The Supplementary Tables S6-10 present detailed results for each information source separately.

**Table 2.** Test for enrichment of diabetes based on text mining/experiment/knowledge base/GWAS catalog for each T2D top ranked pathway.

| Disease |  |  |  |  |  |  | KEGG Pathways |  |  |  |  |  |
| --- | --- | --- | --- | --- | --- | --- | --- | --- | --- | --- | --- | --- |
|  | KEGG_S<br>YSTEMIC<br>LUPUS<br>ERYTHE<br>MATOSU<br>S | KEGG_T<br>YPE_I_DI<br>ABETES<br>MELLITU<br>S | KEGG_A<br>NTIGEN_<br>PROCESS<br>ING_AND<br>PRESEN<br>TATION | KEGG_T<br>YPE_II_D<br>IABETES<br>MELLIT<br>US | KEGG_A<br>XON_GUI<br>DANCE | KEGG_A<br>BC_TRAN<br>SPORTER<br>S | KEGG_ST<br>EROID_H<br>ORMONE<br>BIOSYN<br>THESIS | KEGG_M<br>ATURITY<br>ONSET_<br>DIABETE<br>S_OF_TH<br>E_YOUN<br>G | KEGG_M<br>APK_SIG<br>NALING_<br>PATHWA<br>Y | KEGG_P<br>URINE_M<br>ETABOLI<br>SM | KEGG_SP<br>LICEOSO<br>ME | KEGG_C<br>ELL_CYC<br>LE |
| Central diabetes insipidus | 0,23031816 | 9,8171E-06 | 4,4409E-16 | 0,05458482 | 0,08795732 | 0,56471018 | 0,64653992 | 0,07894318 | 9,81734E-10 | 0,573959154 | 0,908099084 | 1 |
| Diabetes | 2,9661E-08 | 2,341E-07 | 1,1124E-13 | 3,6274E-08 | 1,9207E-14 | 7,648E-08 | 1,2627E-09 | 9,0996E-05 | 0 | 1,24662E-07 | 0,53303145 | 8,47748E-08 |
| Diabetes insipidus | 0,00339478 | 2,8241E-11 | 1,7875E-14 | 1,6241E-07 | 0,01323446 | 0,10560936 | 0,2063247 | 1,5321E-14 | 1,50954E-09 | 0,000783502 | 0,99500288 | 0,265715879 |
| Diabetes mellitus | 1,2517E-07 | 1,5687E-10 | 0 | 9,9522E-12 | 2,4425E-15 | 1,6363E-08 | 6,9389E-14 | 1,0593E-06 | 0 | 2,16494E-05 | 0,93063863 | 2,19911E-06 |
| Dipsogenic diabetes insipidus | 1 | 1 | 1 | 1 | 1 | 1 | 1 | 1 | 0,001299357 | 1 | 1 | 1 |
| Latent autoimmune diabetes in adults | 2,0255E-10 | 0 | 0 | 3,6844E-12 | 1 | 0,49706502 | 0,57657132 | 0 | 0,052808884 | 0,917196321 | 0,860888458 | 0,565832567 |
| Lipoatrophic diabetes mellitus | 0,67012054 | 2,3064E-05 | 1 | 0 | 1 | 0,31039468 | 0,37166429 | 1,9913E-11 | 0,071337175 | 0,740001354 | 1 | 1 |
| Nephrogenic diabetes insipidus | 0,99045573 | 0,16998362 | 9,77E-05 | 0,00016583 | 0,00451679 | 0,00395592 | 0,12374924 | 0,01023961 | 1,56861E-07 | 1,56628E-07 | 0,988594491 | 0,41919612 |
| Nephrogenic diabetes insipidus type 2 | 1 | 1 | 1 | 1 | 1 | 1 | 1 | 1 | 1 | 0,20383297 | 1 | 1 |
| Neurohypophyseal diabetes insipidus | 0,69672354 | 0,00600583 | 0,00079024 | 0,06485783 | 1 | 0,32955368 | 0,39341522 | 0,0212469 | 0,425198371 | 0,765244297 | 1 | 1 |
| **Pancreatic hypoplasia-diabetes-congenital heart disease syndrome** | 1 | 0,06099868 | 1 | 6,9372E-07 | 1 | 0,06531809 | 1 | 3,0544E-10 | 1 | 1 | 1 | 1 |
| **Permanent neonatal diabetes mellitus** | 1 | 0,00341541 | 0,70750794 | 0 | 1 | 0,1442738 | 0,57026734 | 0 | 0,000207584 | 0,034991354 | 1 | 0,846923326 |
| **Prediabetes syndrome** | 0,00352369 | 4,814E-12 | 2,7832E-10 | 0 | 1,5749E-05 | 0,00467562 | 0 | 4,4409E-16 | 0 | 0,175361285 | 0,999670115 | 9,18764E-05 |
| **Transient neonatal diabetes mellitus** | 1,1382E-06 | 6,6232E-07 | 0,05180899 | 6,8882E-11 | 0,83109246 | 0,09931169 | 0,40630118 | 0 | 0,655878504 | 0,913237424 | 0,831092458 | 0,371230641 |
| **Type 2 diabetes mellitus** | 0,0012246 | 1,7838E-10 | 1,2546E-14 | 1,9673E-13 | 1,1102E-15 | 6,7198E-08 | 6,6613E-16 | 1,2587E-07 | 0 | 0,000165776 | 0,991259268 | 3,18823E-08 |
| **X-linked nephrogenic diabetes insipidus** | 1 | 1 | 0,00637804 | 1 | 0,11383984 | 0,13553874 | 1 | 1 | 0,015724755 | 0,002408965 | 0,280751867 | 0,835459666 |

Values show the P-value obtained from the enrichment test.

#### Core genes in the most significantly associated pathways

In our investigation of pathways highly associated with T2D, we focused on genes within the top-ranked pathways, identified with a gene-level *P*-value less than 5×10^−8^ (Figure 7A). Highly associated pathways such as KEGG “*Type I Diabetes*”, “*Antigen processing and presentation*”, and “*Systemic lupus erythematosus*” share several genes, particularly HLA class I and II paralogs (*HLA-DRB1*, *HLA-DQB1*, *HLA-DQA1*, *HLA-B*). Both class I and II molecules play important roles in the immune system, including antigen presentation to T cells and regulation of immune response (44). Additionally, genes such as *LTA* and *TNF* from the tumor necrosis factor family were also associated with these pathways. *LTA* and *TNF* encode multifunctional proinflammatory cytokines, contributing to regulating diverse biological processes, including cell proliferation, differentiation, apoptosis, lipid metabolism, and coagulation (45, 46). Importantly, all these genes play roles in inflammatory and immunostimulatory responses.

**Fig. 7.**
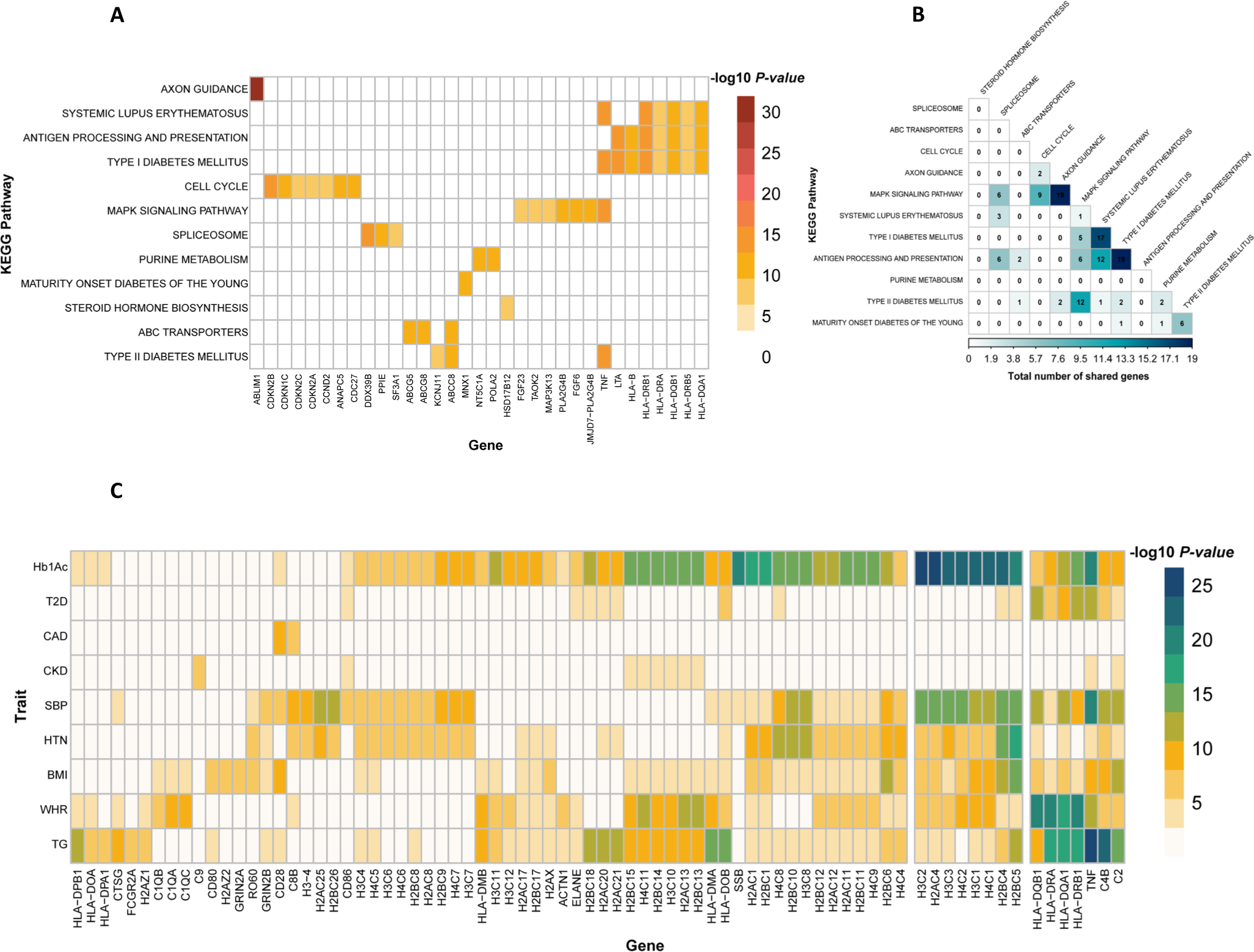
Core genes in the most significantly associated pathways. **A.** Genes with high association (gene-level P-value < 5×10−8) within top-ranked pathways for T2D, with zero values indicating absence in the respective pathway. **B.** Overlapped genes in leading T2D pathways, denoting the count of shared genes between two pathways. **C.** Genes highly associated (gene-level P-value < 5×10−8) in the KEGG pathway “Systemic lupus erythematosus” for each trait.

To explore whether certain genes consistently contribute to disease associations at the pathway level, we selected the “*Systemic lupus erythematosus*” pathway as an exemplar, given its significant association with all examined traits. This pathway encompasses a total of 102 genes. We identified 66 genes within this pathway with gene-based significance (gene-level *P*-value < 5×10^−8^) in at least one trait (Figure 7C). Notably, eight genes from this pathway were found to be associated with at least five traits, showcasing their potential as key contributors. These genes include TNF, HLA class I and II paralogs (*HLA-DRB1*, *HLA-DQB1*, *HLA-DQA*), genes functioning in the classical pathway of the complement system (*C4B*), and *(H2BC5*, *H3C1*, and *H4C1*), all of which have known implications in immunological responses and inflammatory processes (45–49).

## DISCUSSION

The aim of the current study was to propose a novel gene set prioritization approach using single and multiple-trait BLR models. The objectives were not only to identify gene sets associated with individual traits but also to elucidate shared genetic components among different phenotypes. By examining core genes within prioritized pathways, we aimed to enhance biological interpretation, leading to a broad understanding of the genetic landscape governing human complex traits. Our model proved highly effective, as evidenced by extensive simulations and application to T2D and eight related traits. The findings of this study provided valuable insights into the biological mechanisms underlying the studied traits. Further research based on these insights could potentially lead to the identification of promising drug targets for future investigation and therapeutic intervention.

The simulation study provided valuable insights into the performance and robustness of the BLR model. The impact of various gene set specific factors, such as gene set size, the number of causal genes, and their proportion within the gene set, was evaluated in simulated gene sets. One notable finding was the positive effect of increasing the number of causal genes on the F1 score, suggesting that the cumulative effect of more causal genes contributes to a stronger signal, facilitating the BLR model’s ability to distinguish true associations. Conversely, enlarging gene sets with an equal number of causal genes tended to decrease the F1 score, possibly due to a dilution effect where additional non-causal genes in larger gene sets contribute to decreased performance.

The trait-specific factors such as heritability (h²=30% or 10%), proportion of causal variants (π=0.001 or 0.1), genetic architecture (GA1 and GA2), and disease prevalence (5% and 15%), were chosen to mirror real-world scenarios and capture the complexity of different traits genetic architecture. As expected, our model performed significantly better, as evidenced by a higher F1 score, for simulated phenotypes with a lower proportion of causal variants. This improvement suggests that the BLR model can more effectively discern genuine associations from background noise in scenarios with a limited set of causal variants, leading to enhanced detection of true signals.

Similarly, a higher F1 score was observed for scenarios with higher heritability (h²). Elevated heritability implies a stronger genetic influence on the trait, rendering it more amenable to genetic modelling. Consequently, the model’s ability to accurately identify associated gene sets was enhanced when genetic factors substantially influence trait variation. For simulated phenotypes characterized by a few SNPs with large effect sizes (GA1), the model consistently outperformed scenarios with a more complex genetic architecture (GA2). This aligns with the notion that larger effect sizes contribute to a stronger and more discernible genetic signal, enhancing the model’s precision in identifying significant associations within gene sets. Our model exhibited enhanced performance in binary simulated phenotypes with higher prevalence. A higher prevalence indicates a larger proportion of affected individuals, providing more informative data for the model to identify true associations. The increased prevalence amplifies the genetic signals, aiding the model in more accurately prioritizing gene sets associated with the trait. The observed performance of our model across various trait-specific factors validates its effectiveness. It aligns with our expectations, suggesting its potential utility in deciphering genetic associations and prioritizing relevant gene sets.

In both single-trait and multiple-trait BLR analyses of real GWAS summary data, the pathway “*Type II diabetes mellitus*” emerged as a robustly associated pathway with T2D, underscoring its essential role in the pathogenesis of the disease. This pathway is integral to various key processes involved in T2D development, including insulin signalling, regulation of glucose uptake, and metabolism (50, 51). Among the key genes associated with T2D within this pathway are *KCNJ11* (Potassium Voltage-Gated Channel Subfamily J Member 11) and *ABCC8* (ATP-Binding Cassette Subfamily C Member 8), both of which interact with the ATP-sensitive potassium channel. *KCNJ11* and *ABCC8* play crucial roles in maintaining glucose homeostasis, primarily by regulating insulin secretion and glucose metabolism. Dysregulation of these genes disrupts the delicate balance of glucose levels, contributing to the hyperglycaemia observed in T2D (52, 53). Notably, *KCNJ11* and *ABCC8* are targets for commonly prescribed blood glucose-lowering medications, highlighting their clinical relevance in T2D management and emphasizing the therapeutic potential of interventions targeting these pathways (54, 55).

The “*Type I Diabetes Mellitus*” pathway exhibited a strong association with T2D despite this pathways primarily focus on molecular and cellular processes specific to type 1 diabetes (56). This intriguing finding suggests the presence of potential shared mechanisms or specific genes within the Type 1 Diabetes pathway that may interact with or influence the molecular pathways underlying T2D. For instance, several genes within this pathway are associated with the MHC class II locus, a region implicated in immune-mediated processes. Emerging evidence suggests that the genetic architecture of type 1 and type 2 diabetes may harbour common components within the HLA class II locus (47).

Furthermore, the identification of the “*Maturity onset diabetes of the young*” (MODY) pathway adds another layer of complexity to our understanding of T2D. MODY represents a specific monogenic form of diabetes, accounting for approximately 2% of European individuals with T2D (57). While traditionally considered distinct entities, recent studies have shed light on potential connections between MODY and T2D pathogenesis. Emerging evidence suggests that dysregulation of MODY pathways may adversely impact islet function, leading to impaired insulin secretion and glucose metabolism, thereby contributing to the development of T2D (58, 59).

Pathways such as KEGG “*Type I Diabetes*”, “*Antigen processing and presentation*”, and “*Systemic lupus erythematosus*” shared several genes associated with T2D. Remarkably, these genes are vital components of the immune system, playing crucial roles in immune responses. The presence of these immune-related genes within T2D-associated pathways underscores the significance of immune dysregulation in T2D pathogenesis. Indeed, mounting evidence has established a compelling link between chronic low-grade, highlighting inflammation as a key driver of T2D development and progression (60–62).

The application of the BLR model to real data yielded robust insights into known pathways associated with the investigated traits. Our analyses revealed that the multiple-trait analysis consistently outperformed the single-trait analysis across all traits, effectively identifying more pathways. This enhanced performance was attributed to the increased statistical power of the multiple-trait analysis in detecting pathways associated with the trait of interest. Notably, pathways identified through the multiple-trait analysis exhibited higher PIP values, indicating greater significance, and reinforcing that integrating information from multiple traits enhances the detection of shared genetic factors underlying complex traits. These findings support our initial hypothesis and underscore the utility of the BLR model in elucidating the genetic architecture of multifactorial traits.

Understanding the genetic factors behind complex traits can provide valuable insights into the pathogenesis of diseases. Our approach can discover potential drug targets and personalized therapeutic interventions by identifying the interplay between genetic variations and biological pathways. We have validated our findings through enrichment analysis using diverse public resources, ensuring the reliability and robustness of our results. This further supports the translational potential of our findings for clinical and therapeutic applications.

Our BLR modelling strategy has several advantages: First, BLR models utilize external GWAS summary data and LD reference data. They account for LD and can handle different types of genomic features, including gene regions, regulatory feature regions, and other genomic features. These models combine summary statistics from various sources, making them flexible and versatile tools that extend the utility of gene set analysis in genomics. Second, the multiple-trait Bayesian BLR model introduces a novel approach to gene set analysis, specifically designed to explore the associations between gene sets and multiple correlated traits. The model efficiently identifies gene sets relevant across different traits by performing regularization and variable selection concurrently. Moreover, it enables the utilization of information from correlated traits, genders, and ancestries, facilitating a cross-trait analysis approach. This method aims to deepen our understanding of the genetic foundations of human traits, promoting a more comprehensive examination of genetic data across diverse study populations. Third, the BLR models simultaneously perform regularization and variable selection, enabling them to handle a larger number of gene sets and thereby enhancing their analytical and interpretative potential compared to standard MAGMA. Fourth, the BLR models facilitate the fitting of multiple gene set categories, enabling the models to manage more gene sets and contribute differently to the trait.

Our study has certain limitations that need to be considered. One of these constraints is our reliance on widely used pathway resources such as KEGG, which inherently have limitations. These resources may lack high resolution in defining biological pathways and contain a limited number of genes compared to genome-wide datasets. Additionally, they tend to prioritize well-known pathways while potentially overlooking fewer common ones. However, despite these limitations, the KEGG database remains a valuable resource for gaining insights into cellular processes and molecular interactions. The lack of tissue and cell specificity further adds to potential biases in our analysis, constraining our findings within these limitations. Another aspect of our approach is that our pathway-based analysis focuses on genetic variants within gene regions, overlooking a significant number of variants in non-coding regions. This limitation results in information loss for non-coding variants or genes without assigned pathway information, limiting the scope of our analysis in capturing the entire genetic landscape. Moreover, the pathways identified and prioritized by our BLR model are inherently tied to the genetic variants catalogued in the GWAS, potentially overlooking crucial biological insights if specific relevant variants are not included or adequately represented in the GWAS data. Despite these constraints, our study provides valuable insights into the potential of pathway-based analyses in unravelling the underlying mechanisms of complex diseases.

In conclusion, our study introduces a novel approach for prioritizing gene sets using single and multiple-trait BLR models. Through extensive simulations and analyses of real traits, we have demonstrated the efficacy of the BLR model in prioritizing pathways for complex traits. The multiple-trait BLR model, in particular, stands out as a flexible framework capable of uncovering shared genetic pathways and highlighting the interconnected nature of trait genetics. Our approach paves the way for advancements in genomics, systems biology, and personalized medicine by identifying relevant pathways associated with complex traits. While our findings showcase the promise of the BLR model, further research is needed to address potential limitations and broaden its applicability in diverse research settings.

## DATA AVAILABILITY

The genetic and phenotypic data utilized in our study were obtained from the UK Biobank Resource under Application Number 96479, https://bbams.ndph.ox.ac.uk/ams/. The datasets analyzed during the current study are available for download from the following URLs: Type 2 Diabetes KP genetic association datasets, https://t2d.hugeamp.org/datasets.html; DISEASES: Disease-gene associations mined from literature, https://diseases.jensenlab.org. 1000 Genomes Project reference datasets were downloaded from the Centre for Neurogenomics and Cognitive Research (CNCR) website, including g1000_eur.zip, g1000_eas.zip, and g1000_sas.zip for the respective populations, https://ctg.cncr.nl/software/MAGMA/ref_data/. Ensembl gene annotations were obtained from: ftp.ensembl.org/pub/grch37/current/gtf/homo_sapiens/Homo_sapiens.GRCh37.87.gtf.gz

## CODE AVAILABILITY

The BLR prioritization approach is available as a part of an open-source R package at https://github.com/psoerensen/gact.

## AUTHOR CONTRIBUTION

P.S. conceived the study; M.S., Z.B., and T.G. pre-processed the data; T.G. performed all analyses, with conceptual input from P.S., P.D.R., A.J.H. and M.F.K; T.G. and P.S. drafted the manuscript; all authors contributed to and approved the final manuscript.

## ETHICAL APPROVAL

Human studies in the UK Biobank project have received approval from the Ethics and Governance Framework (EGF), which ensures data and sample usage adheres to scientific and ethical standards. The consent to participation will apply throughout the lifetime of the UK Biobank, unless participants withdraw, and involves the collection and storage of biological samples (blood, saliva, urine) and electronic health records (GP, hospitals, dental, prescriptions). Individual data is anonymized, with each research project receiving its own anonymized dataset. The ethics committee waived the need for written informed consent.

