## Supplementory Figure 1 for "Evaluation of Bayesian Linear Regression Models for Gene Set Prioritization in Complex Diseases"

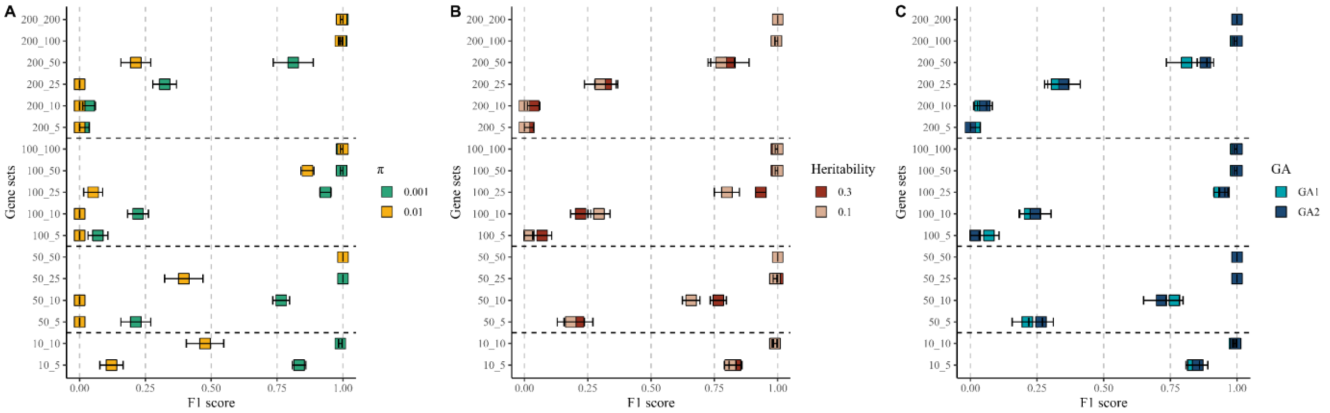


**Fig. S1**. **Evaluation of BLR model performance in simulation scenarios (quantitative traits)**. Scenarios were compared by varying a specific property while keeping others constant. **A.** Illustrates the impact of varying the proportion of causal markers (π). **B.** Demonstrates scenarios with varying heritability ($h^2$). **C.** Compares two genetic architecture scenarios, GA1 and GA2. The y-axis represents gene sets, with the first number indicating the size of the gene sets and the second number representing the number of causal genes within the gene set. The x-axis displays the $F_1$ score. Points represent mean values across 10 replicates, and error bars indicate standard errors.
